## Supplementary Information for "PandoGen: Generating complete instances of future SARS-CoV-2 sequences using Deep Learning"

### PandoGen: Supplementary Document

#### 1 Algorithm for sequence comparisons with ambiguous characters

The algorithm for comparing ambiguous characters to unambiguous characters is shown in Algorithm 1.

**Algorithm 1** Sequence Comparison between sequences X, Y

```
if length(X) != length(Y) then
  return "Unequal"
else
  for i in 0 to length(X) - 1 do
    if X[i] != Y[i] then
      if X[i] == "X" or Y[i] == "X" then
        continue loop
      end if
      if X[i] == "B" and Y[i] ∈ {"D", "N"} then
        continue loop
      end if
      if X[i] == "J" and Y[i] ∈ {"L", "I"} then
        continue loop
      end if
      if X[i] == "Z" and Y[i] ∈ {"Q", "E"} then
        continue loop
      end if
      if Y[i] == "B" and X[i] ∈ {"D", "N"} then
        continue loop
      end if
      if Y[i] == "J" and X[i] ∈ {"L", "I"} then
        continue loop
      end if
      if Y[i] == "Z" and X[i] ∈ {"Q", "E"} then
        continue loop
      end if
      return "Unequal"
    end if
  end for
  return "Equal"
end if
```

#### 2 Details on scaling PandoGen sample size

We scaled the sample output size of PandoGen to see what fraction of the sequence space succeeding the training period, can PandoGen capture. To prepare these comparisons, we selected the sampling configurations for PandoGen that results in the highest number of forecasted sequences from experiments in Main document Figure 3. For comparison, we selected a sampling configuration of "Prot GPT2 unenumerated" with the highest number of forecasted sequences as well.

We ran three sampling experiments with these configurations generating 16,384 sequences in each sampling run. Figure 1 shows the number of novel lineages forecated by the models over time. PandoGen forecasts tens of novel lineages, whereas Prot GPT2 forecasts virtually no new lineages. The forecasted lineages from PandoGen saturate after week 13. The figure also shows the number of novel lineages from the methods as a fraction of all novel lineages first reported in GISAID in the same time periods. Significantly, after the end of the training period, until the 10th week after the training period, PandoGen forecasts close to 25% of all lineages first reported in GISAID in that same period.

#### 3 Performing Pango lineage reassignments

We downloaded the full SARS-CoV-2 genomic sequences as a fasta file from GISAID on 2023-10-16. We also downloaded the Pangolin release v4.3.1 (`https://github.com/cov-lineages/pangolin/releases/tag/v4.3.1`). We created a script run_pangolin_batched.py as a manager script to run Pangolin on multiple nodes in a batched fashion. We ran lineage assignment in batches because the tool crashes when trying to perform assignment for some sequences, hence batched launch allows us to isolate the failing cases.

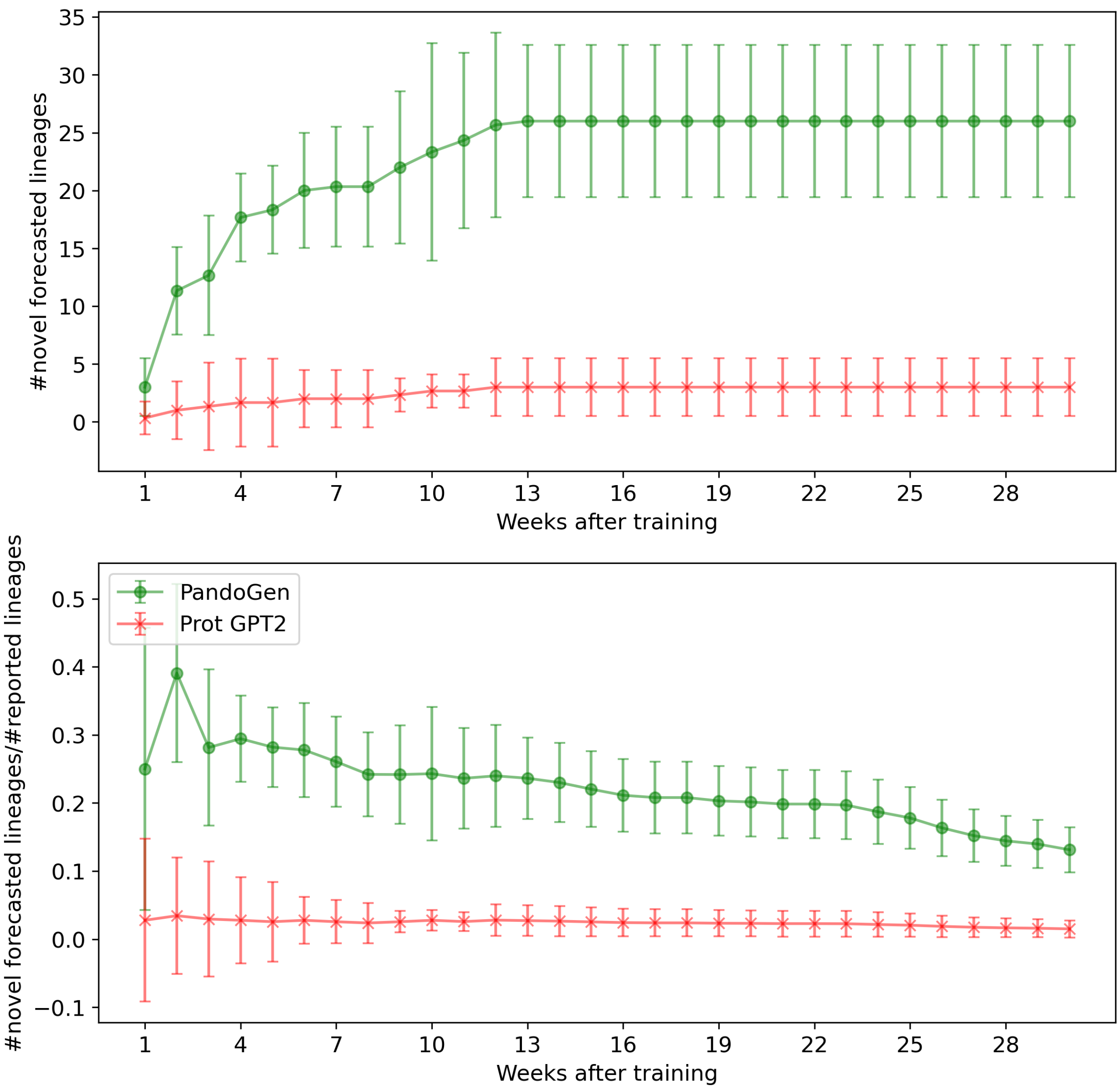


Figure 1: Lineage forecast efficacy. Above: The number of lineages predicted until a given week after the end of the training period. Below: novel lineages forecasted as a fraction of all novel lineages reported in GISAID in the same time period. The spreads represent 95% C.I.

Our experiments in the main article are based on GISAID source data downloaded on 2022-12-02. For each accession in this file, we compared the lineage assignment in the source file, and the new lineage assignment created by us. The following are the results.

- Number of accessions in GISAID source file on which our experiments are based: 14,035,944
- Number of unique Spike sequences in GISAID source file: 852,529
- Number of accessions in GISAID source file without a lineage assignment: 506
- Number of deduplicated accessions in GISAID fasta file: 16,104,063
- Number of accessions in GISAID fasta for which lineage assignment did not work: 20
- Number of accessions for which lineage assignments mis-match for the two cases: 613,093
- Number of unique Spike sequences for which lineage assignments mis-match for the two cases: 112,116
- Number of accessions in GISAID source file for the training period: 1,992,335
- Number of unique Spike sequences in GISAID source file for the training period: 104,106
- Number of accessions in the training period with lineage mismatch: 43,142
- Number of unique Spike sequences in the training period with lineage mismatch: 9,837
